# Ignet 2.0 and Vignet: An Ontology-Driven Web Platform for Biomedical Gene Interaction Discovery and Visualization

**DOI:** 10.64898/2026.06.02.729682

**Authors:** Sayed Asaduzzaman, Benu Bansal, Parker Combs, Jie Zhang, Hasin Rehana, Brett McGregor, Yongqun He, Junguk Hur

## Abstract

**Background:** The expansion of biomedical literature demands systematic ontology-guided discovery of gene interactions, vaccine mechanisms, drug associations, and adverse events. Existing platforms such as STRING, DisGeNET, and PubTator fall short of providing a unified, freely accessible system that integrates ontology-based semantic interaction classification, vaccine-focused heterogeneous network construction, and Artificial Intelligence-assisted evidence retrieval.

**Results:** Ignet 2.0 and Vignet are freely accessible dual-platform systems that combine PubMed literature mining, BioBERT-based interaction scoring for millions of gene-gene co-occurrence pairs and integrate three biomedical ontologies and one curated drug resource, Interaction Network Ontology (INO), Vaccine Ontology (VO), Human Disease Ontology (HDO), and DrugBank. Ignet 2.0 supports gene interaction discovery, gene set enrichment retrieval of BioBERT-scored GenePair evidence, and AI-assisted summarization through BioSummarAI. Vignet extends these features with VO-guided Vaccine Exploration, VacPair interaction scoring, and the creation of vaccine, gene, drug, and disease networks in VacNet. A public Representational State Transfer Application Programming Interface (REST API) and Model Context Protocol (MCP) endpoint enable real-time integration, fostering trust in biomedical knowledge discovery.

**Conclusion:** Ignet 2.0 and Vignet are scalable, ontology-guided biomedical knowledge platforms that facilitate evidence-based gene interaction analysis, vaccine-focused semantic exploration, and AI-assisted knowledge discovery. Their real-time PubMed data integration ensures up-to-date insights; however, users should consider validation processes and potential lags in incorporating the latest experimental data, which may affect the reliability of immediate data.

**Availability:** Ignet 2.0: https://ignet.org/ignet; Vignet: https://ignet.org/vignet/

## 1. Background

The rapid increase in biomedical literature makes research on vaccine and drug ontologies challenging. PubMed lists over 40M citations and adds millions more each year [1]. Gene-level mechanisms for vaccine immune responses, drug interactions, and adverse events are scattered across thousands of publications. Conventional keyword search returns only an article list, not structured knowledge. For example, if we search for “Influenza Vaccine” in PubMed, it returns thousands of abstracts. However, it does not show the most co-occurring genes, drugs, and diseases, along with their interaction types and supporting evidence. This gap between the amount of literature and discoverable knowledge is a major problem that an ontology-guided mining process aims to address in the current research world [2,3]. Recently, several text mining platforms have emerged to address the issue. PubTator3 is such a platform that performs automated recognition and annotation of gene, disease, chemical, and species in the literature [4]. A widely used biological database and web resource in molecular biology is STRING[5], which connects, validates, and predicts protein interactions [6]. On the other hand, DisGeNET [7] collects gene-disease associations from curated and mined sources [8]. Although various text-mining and database tools are available, a unified framework for the systematic retrieval and analysis of vaccines, genes, drugs, and diseases within a single ontology-based system is lacking.

The Comparative Toxicogenomic Database (CTD) updated its data through manual curation focused on chemicals, providing current and complete information. However, CTD does not classify interaction types using ontologies or validate evidence with machine learning. This lacks vaccine-specific ontology integration for class-level queries and cannot be updated daily with new PubMed content. Therefore, none of the above-mentioned platforms provides a cross-entity network that integrates vaccines, genes, drugs, and diseases within a single ontology-based visualization, which is much needed for vaccine and drug ontology research [9].

Ontology-based literature mining improves the discovery of biomedical interactions by integrating gene normalization, semantic interaction classification, and hierarchical ontology reasoning. Frameworks such as SciMiner enable ontology-guided enrichment analysis and improve literature retrieval. Such as, VO hierarchy traversal increased vaccine-related PubMed retrieval by approximately 12% compared with keyword-only searches [3,10]. INO interaction type classification and four centrality metrics (degree, eigenvector, closeness, and betweenness) were used to rank gene importance. This was done in Ignet version 1, which first provided centrality- and ontology-based interaction classification for literature-mining gene networks [11].

Transformer-based biomedical language models have created new opportunities for ontology-based mining. BioBERT, trained on PubMed abstracts and articles, outperforms general language models for extracting biomedical relations. It can score sentence-level interactions more accurately than keyword co-occurrence. Large language models (LLMs) such as GPT-4o provide advanced summarization and question-answering capabilities [12]. AI-powered synthesis, Ignet 2.0, and Vignet operate as an open-access dual-platform system. This system integrates three biomedical ontologies and one curated drug resource: VO, INO, HDO, and DrugBank, and analyzes new PubMed abstracts daily. It uses SciMiner-based entity recognition and BioBERT-based confidence scoring for gene-gene interaction evidence in millions of records. For literature summaries, BioSummarAI creates concise, evidence-based overviews linked to supporting PubMed citations and PMIDs; gene-pair evidence includes BioBERT confidence scores. The platform can also be accessed directly by AI assistants through a public MCP endpoint.

## 2. Implementation

Ignet 2.0 and Vignet are implemented as a unified dual-platform system that shares a common backend infrastructure, natural language processing (NLP) pipeline, and ontology integration layer. The subsequent subsections outline the system architecture, the literature mining pipeline, the database design, the integrated ontologies, the principal platforms, and API interoperability.

### 2.1 System Overview

Figure 1 shows the architecture of Ignet 2.0 and Vignet, highlighting the overall pipeline, including literature acquisition, daily ingestion (update), NLP-based entity recognition, ontology integration, knowledge base construction, and application services. Articles from the PubMed collection are automatically processed using eXtensible Markup Language (XML) parsing and sentence segmentation pipelines in SciMiner upstream. SciMiner NLP module identifies genes, diseases, drugs, vaccines, and interaction terms, while BioBERT provides interaction confidence scores. Extracted entities are integrated using ontologies such as INO, VO, HDO, and DrugBank. The knowledge base stores gene interactions supported by literature, ontology annotations, and ranked evidence. These resources enable applications such as interactive network visualization, pairwise interaction analysis, enrichment analysis, AI-assisted summarization, REST API services, and MCP-compatible interoperability.

**Figure 1.**
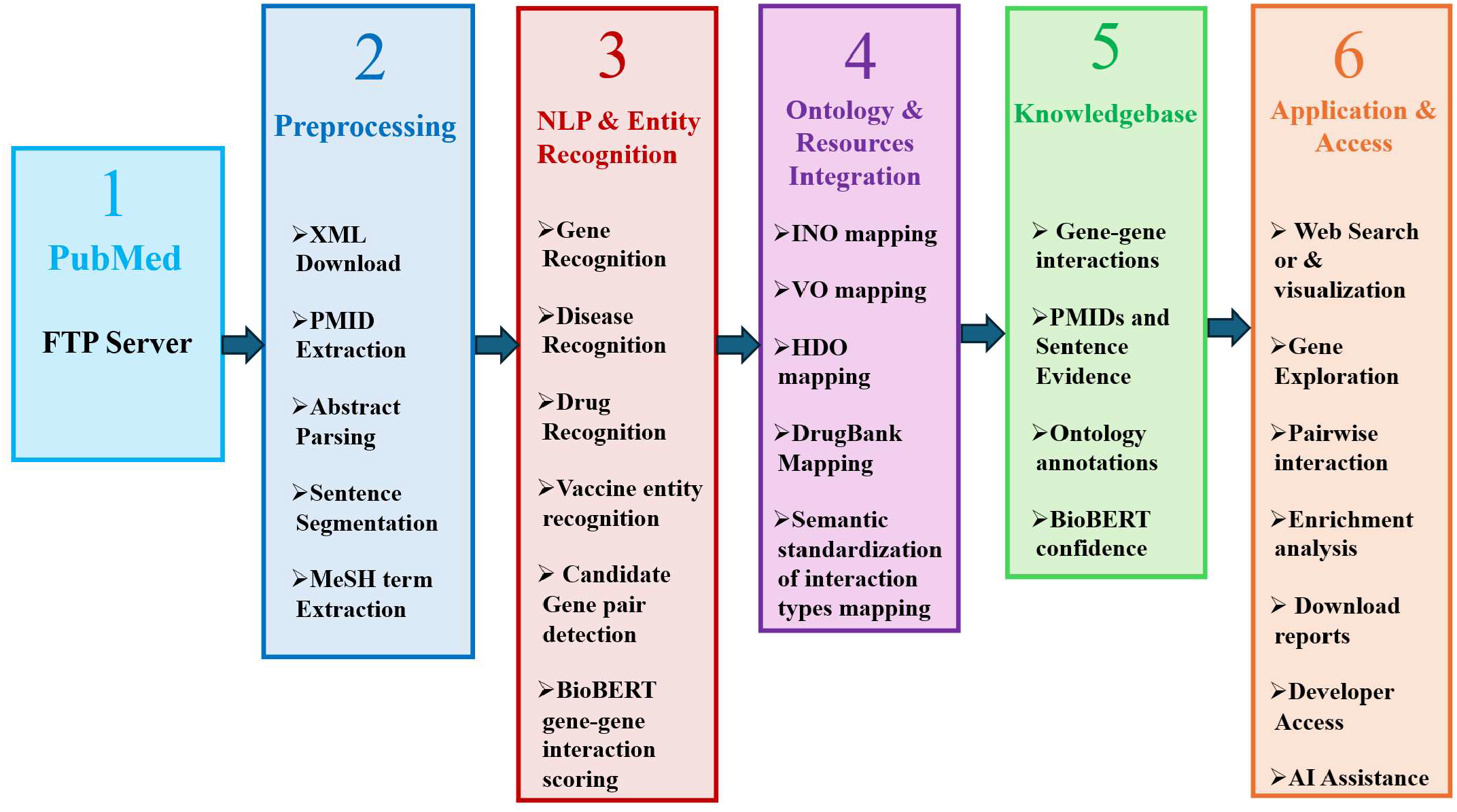
Overview of the System Architecture and Pipeline Flowchart. The workflow consists of six stages: (1) PubMed literature acquisition, (2) preprocessing and sentence extraction, (3) NLP-based entity recognition and BioBERT interaction scoring, (4) ontology and resource integration using INO, VO, HDO, and DrugBank, (5) construction of a biomedical knowledgebase containing gene interactions and supporting evidence, and (6) user-facing applications including network exploration, enrichment analysis, AI-assisted summarization, REST API access, and MCP-enabled interoperability.

### 2.2 Literature Mining and Interaction Extraction

Ignet 2.0 uses a multi-stage natural language processing (NLP) pipeline, built on the SciMiner framework, for literature mining and interaction extraction. In the legacy system Ignet, PubMed XML files were downloaded directly from the National Center for Biotechnology Information (NCBI) File Transfer Protocol (FTP) server and processed with a PHP script. In Ignet 2.0, this step is handled by the upstream SciMiner pipeline, which downloads and processes PubMed XML files and transfers preprocessed output files to the Ignet server for batch loading into MySQL. SciMiner employs a Java-based sentence splitter to divide each abstract into sentence-level records, each linked to its source PMID, before transferring the output to the Ignet database. SciMiner applies ontology- and dictionary-based entity recognition using modules such as Host gene, INO, VO, HDO, and DrugBank. Sentences with multiple recognized entities are scored to identify potential interactions, which are then filtered to remove low-confidence matches. The resulting interactions and supporting sentences are stored in a MySQL database for retrieval and network analysis. A BioBERT-based classification module further refines gene-gene interaction confidence scores, thereby supporting ranked evidence retrieval and downstream analysis within the Ignet platform.

### 2.3 Database Architecture

MySQL is used to store the database, with PubMed articles as the primary source anchor. The overall system architecture of the database is shown in **Figure 2**. Each record links to sentence-level entries generated during preprocessing, ensuring that every extracted interaction is traceable to its supporting evidence. Gene-gene interaction records include co-occurring gene pairs, source PMIDs, sentence identifiers, matched surface forms, contextual flags, and BioBERT-derived confidence scores ranked by retrieval. Four ontology annotation tables, INO, VO, HDO, and DrugBank, use a standardized structure to associate PMIDs with matched ontology terms. This design enables cross-referencing among interactions, immunological processes, vaccine concepts, disease annotations, and drug entities. The system architecture supports efficient evidence retrieval, ontology-based network construction, and web-based visualization. The daily incremental update uses a PMID-based reconciliation strategy to refresh only the affected records, eliminating the need to refresh the entire database.

**Figure 2.**
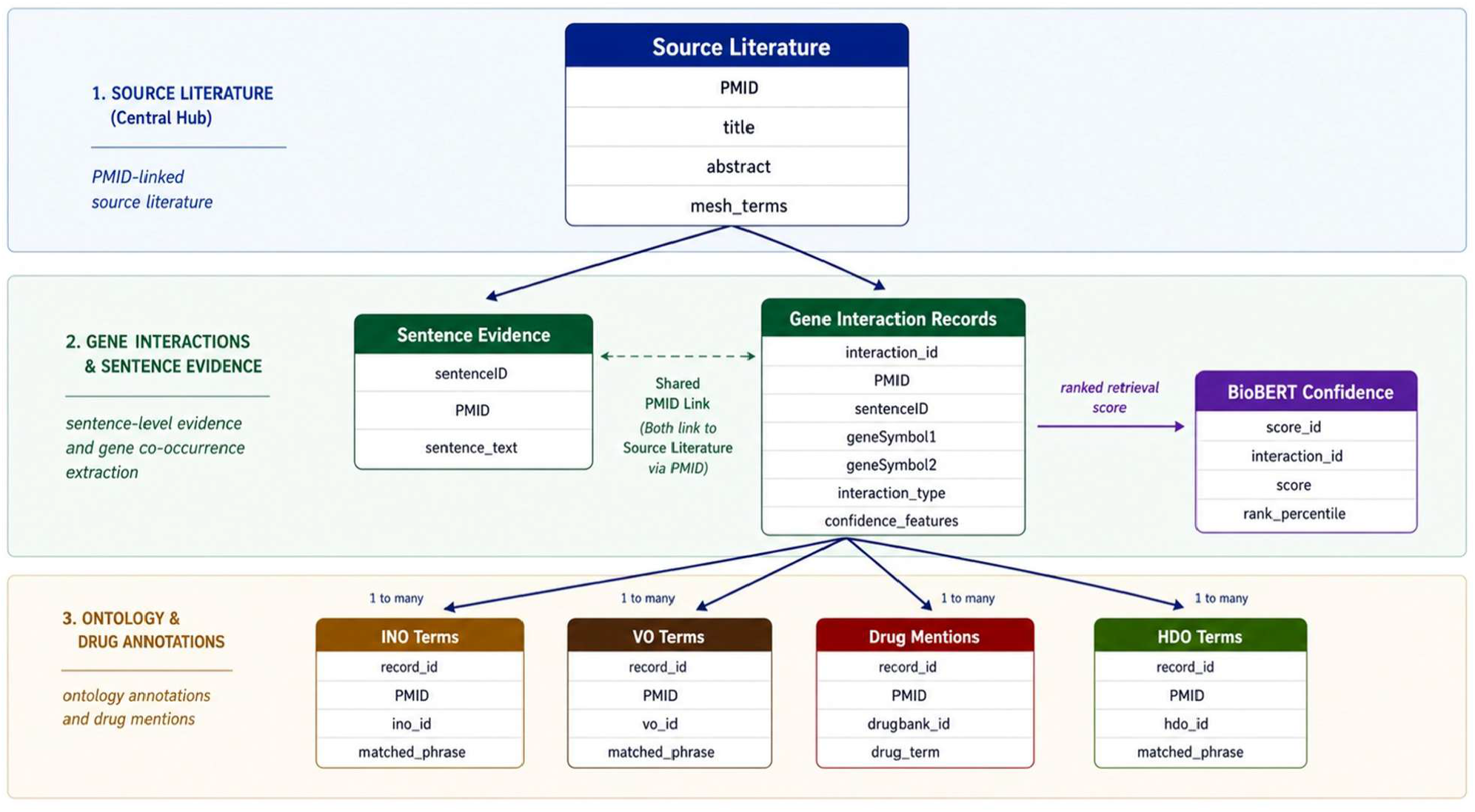
Semantic Database System Architecture. The three-tier schema connects PMID-indexed source literature at the top level to gene interaction records and sentence-level evidence with BioBERT confidence scores at the middle level, and to ontology and drug annotation tables INO, VO, HDO, and DrugBank at the bottom level through one-to-many relationships.

### 2.4 Integrated Ontologies

Ignet 2.0 and Vignet incorporate four biomedical ontologies and curated knowledge resources to facilitate standardized entity normalization, semantic integration, classification, and cross-domain biomedical knowledge interaction, as shown in **Table 1**. The INO provides a controlled vocabulary for classifying gene-gene interaction types extracted from biomedical literature and enables ontology-guided semantic annotation of all international biomedical literature. enabling ontology-guided semantic annotation of all international biomedical literature and enabling ontology-guided semantic annotation of all interaction records in Ignet. VO underpins Vignet’s identification, traversal of the VO hierarchy, and construction of heterogeneous networks in VacNet. The HDO and DrugBank serve as external reference resources for annotating disease and drug nodes in gene report enrichment analyses and in VacNet heterogeneous graphs, thereby enabling the integration of biomedical knowledge across genes, vaccines, diseases, and drugs.

**Table 1.**
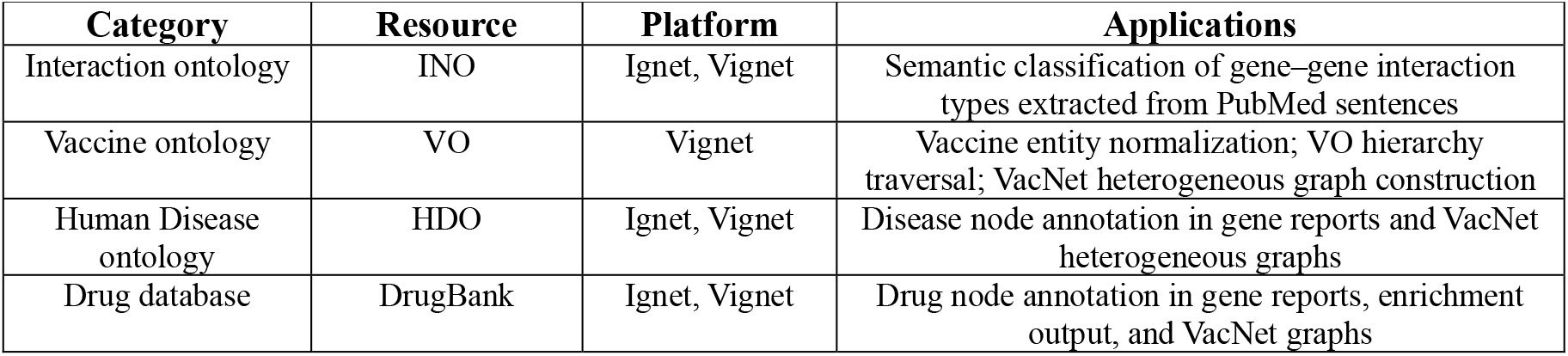
Biomedical Ontologies and Knowledge Resources Integrated in IGNET 2.0 and Vignet.

### 2.5 Key Platform Features

#### 2.5.1 Ignet 2.0

Ignet 2.0 is an integrated platform for biomedical literature mining and interaction analysis, shown in **Figure 3**. It enables large-scale identification of gene co-occurrence and semantic interconnection networks in the PubMed literature. The system combines automated literature mining, BioBERT-based interaction prediction, ontology-guided semantic classification, and interactive exploration tools to support evidence-based biomedical knowledge discovery and translational interaction analysis.

**Figure 3.**
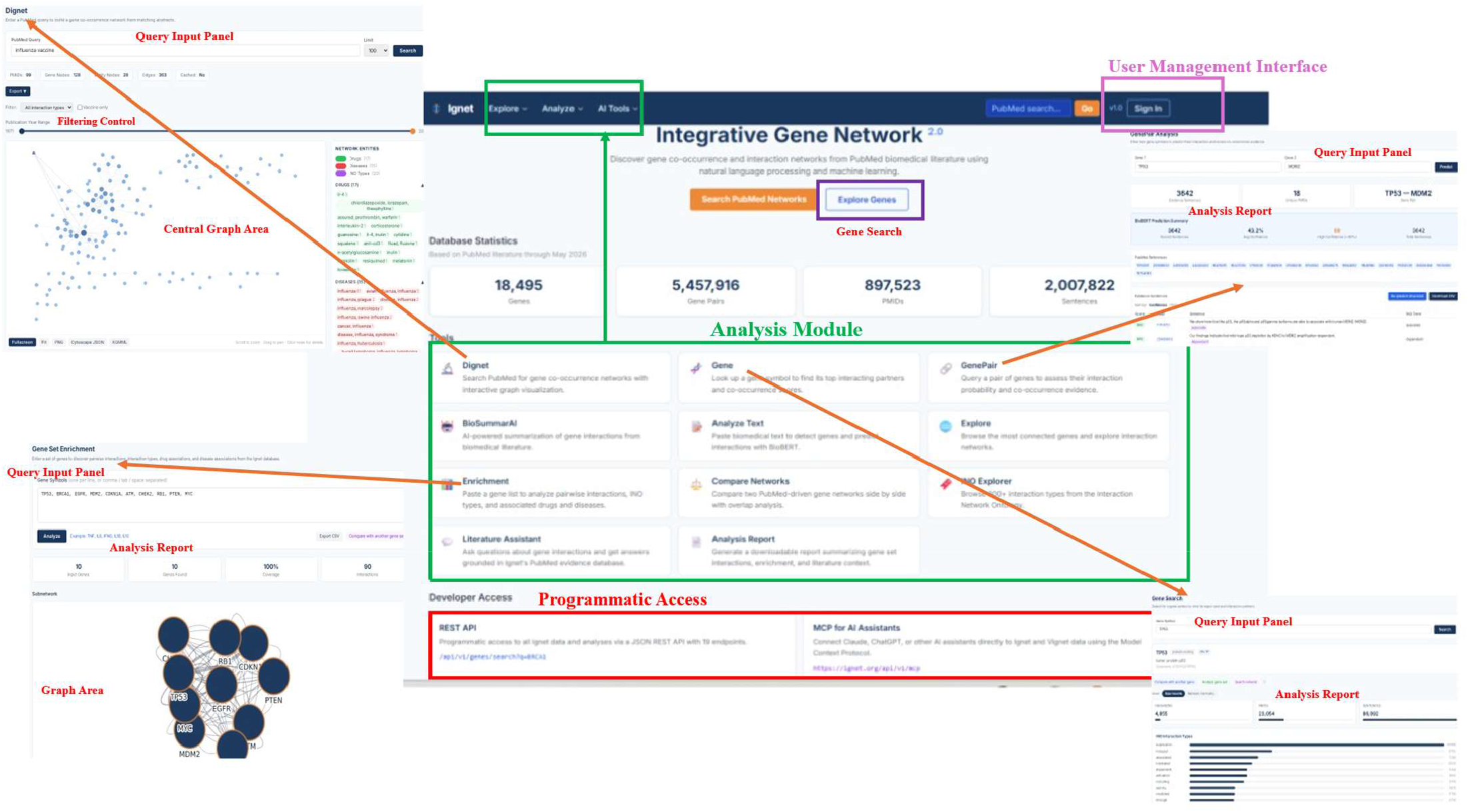
Overview of Ignet 2.0 and its different panels. The figure illustrates the primary components of the Ignet 2.0 platform: Query Input Panels, Filtering Controls, Central Graph Area, Analysis Report sections, User Management Interface, the Analysis Module (which includes tools such as Dignet, Gene, GenePair, BioSummarAI, Analyze Text, Explore, Enrichment, Compare Networks, INO Explorer, Literature Assistant, and Analysis Report), and Programmatic Access options such as REST API and MCP for AI Assistants.

➢ **Gene and Gene Pair Exploration:** The Gene search and Gene Pair modules enable retrieval of interaction partners, PMIDs, INO classified interaction types, drug associations, and centrality rankings. Gene pairs use BioBERT to estimate the probability of interaction between two genes and provide confidence-ranked evidence sentences linked to PubMed references.
➢ **Network Discovery and Exploration:** Dignet builds gene interaction networks from literature by processing PubMed queries with filters for interaction type and entity class. The Compare Networks tool displays two query-driven networks side by side, making it easier to identify unique molecular relationships in different biological contexts.
➢ **Gene Set Analysis and Enrichment:** Users can submit gene lists from RNA-seq or proteomics studies to generate a pairwise interaction subnetwork annotated with INO types, drugs, and diseases. Results are provided as CSV files or downloadable analysis reports to support translational and systems biology research.
➢ **AI-Augmented Information Retrieval:** The Literature Assistant, BioSummarAI, and Analyze Text tools offer PubMed-based question answering, GPT-4o-driven literature summarization, and BioBERT-supported gene interaction, respectively, from user-provided biomedical text.

#### 2.5.2 Vignet

Vignet is a semantic biomedical exploration platform for vaccines, built on the Ignet framework to support ontology-guided knowledge discovery and interaction analysis, as illustrated in **Figure 4**. It features vaccine ontology-based navigation, vaccine-gene association mining, heterogeneous network construction, and AI-assisted evidence interpretation to advance research on vaccine and drug ontologies.

**Figure 4.**
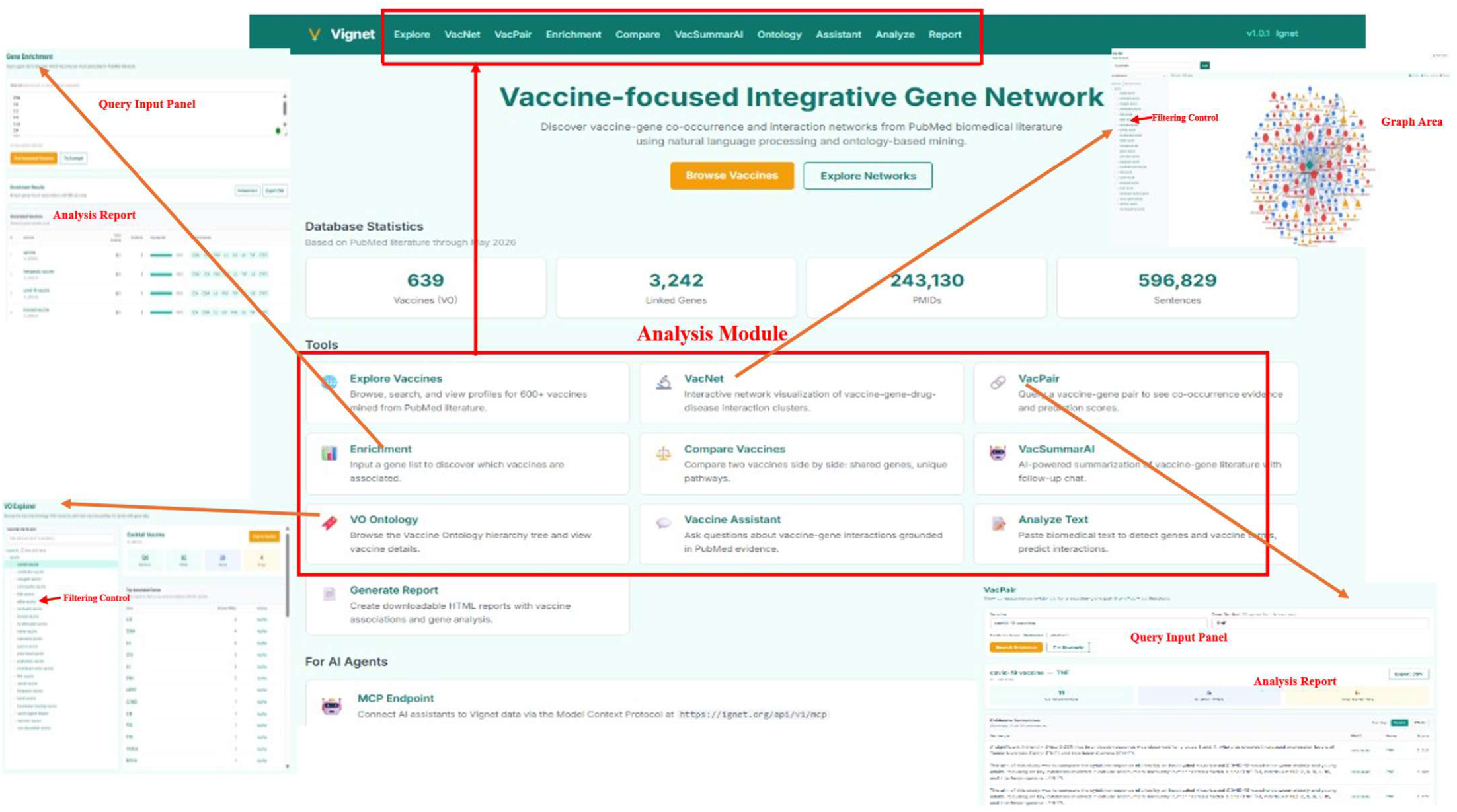
Overview of Vignet and its different panels. The figure illustrates the primary components of the Vignet (Vaccine-focused Integrative Gene Network) platform. These components include the Query Input Panels, Filtering Controls, Graph Area, Analysis Report sections, and the Analysis Module. The Analysis Module comprises tools such as Explore Vaccines, VacNet, VacPair, Enrichment, Compare, VacSummarAI, VO Ontology, Vaccine Assistant, Analyze Text, Generate Report, and MCP Endpoint access for AI Assistants.

➢ **VO Navigation and Discovery:** Vignet supports ontology-guided exploration of vaccines and their subclasses using a VO hierarchy. The platform allows users to navigate vaccine concepts, retrieve ontology-linked categories, and access related biomedical entities and PubMed literature.
➢ **Vaccine Gene Network Construction:** The platform constructs diverse vaccine-gene interaction networks by mining PubMed literature and performing ontology-linked association analysis. It offers interactive visualizations of vaccine-associated genes, diseases, drugs, and classified interaction types.
➢ **Vaccine Gene Comparison and Enrichment Analysis:** Vignet provides comparative and enrichment analysis tools to identify shared genes, interaction patterns, and ontology-linked biological associations across multiple vaccines. These tools support research on immune-response pathways and advance translational vaccine development.
➢ **Vaccine AI Analyzer, Assistance, and Report Generator:** VacSummarAI generates evidence-based literature summaries using GPT-4o. Its integrated AI assists with vaccine-related biomedical questions and automates report generation, incorporating evidence from the literature, interaction networks, and enrichment analyses.

### 2.6 API and MCP Interoperability

Ignet 2.0 and Vignet expose programmatic access via a public REST API (base URL: https://ignet.org/api/v1), and an MCP endpoint (https://ignet.org/api/v1/mcp) is depicted in the interactions. The API provides JSON-based access to gene interaction records, BioBERT-scored gene-pair evidence, PubMed-derived co-occurrence networks, INO ontology terms, gene set enrichment, GPT-4o literature summarization, and VO-guided vaccine–gene associations, as shown in Table 2. Most endpoints have a daily usage cap. The MCP server exposes eight tools spanning both platforms, enabling MCP-compatible AI assistants to retrieve ontology-guided biomedical knowledge and evidence-grounded interaction data in real time without local installation.

**Table 2.**
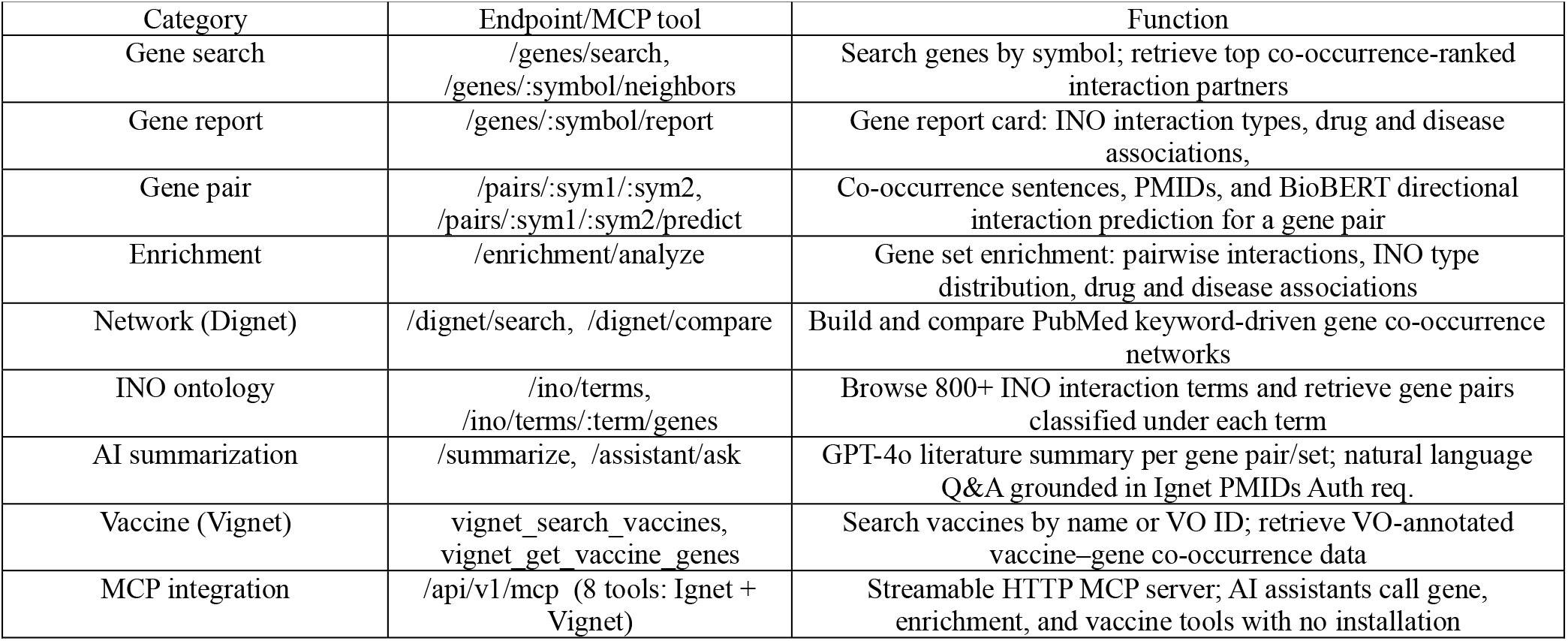
REST API and MCP Blueprint.

## 3 Results

The capabilities of Ignet 2.0 and Vignet were evaluated using representative ontology-guided biomedical discovery workflows that prioritized evidence-based interaction, validation, and exploration of heterogeneous vaccine-centered networks. The following use cases demonstrate the platform’s integration of literature mining, semantic interaction analysis, and AI-assisted biomedical knowledge discovery.

### 3.1 Benchmarking and Comparison

Ignet 2.0 and Vignet offer integrated literature search, ontology-driven analysis, BioBERT-based interaction prediction, and AI-guided exploration of biomedical topics. In contrast to STRING, BioGRID, DisGeNET, MetaScape, KnetMiner, and PubTator, which focus on interaction types, vaccine research, network mapping, or evidence aggregation, as shown in Table 3. Vignet uses the VO to connect vaccines, genes, drugs, and diseases. Together, Ignet 2.0 and Vignet uniquely provide guided interaction analysis, evidence-based literature search, AI-supported interpretation, and exploration of diverse biomedical networks on a single platform.

**Table 3.** Feature Comparison of Similar Tools.

| Feature | Ignet 2.0 | Vignet | STRING<br>[5] | BioGRID<br>[13] | DisGeNET<br>[7] | Metascape<br>[14] | KnetMiner<br>[15] | PubTator<br>[4] |
| --- | --- | --- | --- | --- | --- | --- | --- | --- |
| Literature-mined interactions | Supported | Supported | Partial | No | Partial | Partial | Partial | Supported |
| Ontology integration | Supported | Supported | No | No | Partial | Partial | Supported | Partial |
| Vaccine-centered analysis | Limited | Supported | No | No | No | No | No | No |
| Sentence-level evidence | Supported | Supported | No | No | Limited | No | Limited | Supported |
| BioBERT/AI integration | Supported | Supported | No | No | No | No | No | No |
| Heterogeneous network exploration | Supported | Supported | No | No | Limited | Limited | Supported | No |
| Dynamic PubMed queries | Supported | Supported | No | No | No | No | No | Supported |
| Interactive visualization | Supported | Supported | Supported | Supported | Limited | Supported | Supported | Limited |
| Gene set enrichment | Supported | Supported | No | No | Limited | Supported | Limited | No |
| REST API access | Supported | Supported | Supported | Supported | Supported | No | No | Supported |
| MCP interoperability | Supported | Supported | No | No | No | No | No | No |

### 3.2 Representative Application Workflows

#### 3.2.1 BioBERT-assisted validation of gene and vaccine-associated interactions

Given two biomedical entities, does literature support an interaction between them, and how strong is the evidence?

➢ **System Expectation:** The system retrieves sentences from PubMed, evaluates them using BioBERT, and ranks evidence based on interaction confidence to distinguish mechanistic from incidental co-mentions.
➢ **Needed Modules:** Ignet (GenePair), Vignet (VacPair), BioBERT Scoring, INO annotation modules
➢ **Scenario:** A biomedical researcher investigating immune-response pathways or vaccine-associated signaling mechanisms and wishes to validate whether two genes are functionally associated in the literature. Relying solely on co-occurrence frequency, the researcher uses Gene Pair to retrieve sentence-level evidence, BioBERT confidence scores, and INO-classified interaction types. The workflow enables explainable and evidence-supported interaction types.
➢ **Example:** A user enters TNF and IL6 into the Gene Pair analysis tool, which identified 15,148 evidence sentences from 27 unique PMIDs, with an average BioBERT confidence score of 36.2%. Among these, 12 sentences exhibited high confidence exceeding 80%. When ranked by confidence, the top sentence demonstrated clear mechanistic relationships. When ranked by confidence, the top evidence sentences reflected distinct mechanistic interaction types, including induction (89% confidence), modulation (84% confidence), and binding-related interactions (84% confidence). Applying the same methodology to the COVID-19 vaccine and myeloperoxidase (MPO) using the Vignet VacPair yielded only 2 sentences across 2 PMIDs. Both sentences are clinically significant: a NET-associated MPO biomarker study and a case report of MPO-ANCA positivity following Pfizer-BioNTech COVID-19 vaccination. These results were obtained without manual curation, demonstrating that BioBERT scoring effectively distinguishes mechanistic evidence from incidental co-mentions.

#### 3.2.2 Ontology-guided heterogeneous network exploration across vaccines, genes, drugs, and diseases

How are vaccines, genes, drugs, and diseases connected in a heterogeneous biomedical network?

➢ **System Expectation:** The system enables users to explore direct and indirect relationships between different entity types in a single, ontology-guided graph view.
➢ **Needed Modules:** Vignet (VacNet), Vaccine-gene association, Gene-gene interaction, Cross-entity visualization and filtering, and Drug-disease association modules
➢ **Scenario:** Researchers studying vaccine-associated immune mechanisms seek to clarify connections among vaccine classes, genes, drugs, and diseases using literature-derived semantic relationships. Vaccine classes, genes, drugs, and diseases are the four biomedical entity types unified in the VacNet and VO hierarchy, forming a heterogeneous interaction network that integrates these types within ontology-aware graphs. This system supports both direct vaccine-gene relationships and implicit ontology-based associations. By traversing VO vaccine ontology subclasses, this approach enables cross-domain biomedical knowledge discovery beyond the capabilities of single-entity queries or keyword-based literature searches.
➢ **Example:** A user loads COVID-19 vaccine into Vignet VacNET, generating 189-node, 355-edge network spanning genes (ACE2, IL6, IFNG, TMPRSS2), drugs (remdesivir, Spikevax, dexamethasone), and diseases (diabetes mellitus, lupus, myocarditis), each connected to the vaccine center node via literature-derived edges. Enabling “Show gene -gene interactions” and “Show cross-neighborhood interactions” expands the graph to 855 edges, revealing multi-hop paths linking vaccines to drugs through shared gene intermediaries. Switching to (Combine Vaccine) with implicit child associations produces a 131-node, 166-edge heterogeneous network connecting genes (IL6, IFNG, CD4, CD8A, CTLA4, GZMB), drugs (infliximab, azithromycin, dexamethasone), and diseases (rubella, malaria, diphtheria, hepatitis B, pertussis). The VO hierarchy sidebar highlights data-bearing vaccine classes, guiding researchers to evidence-supported nodes across the full 6796-term VO ontology.

## 5 Discussion

Ignet 2.0 and Vignet address a longstanding challenge in biomedical informatics: the lack of a unified, ontology-driven platform that connects gene interactions, vaccines, diseases, and drugs within a single evidence-based framework. These platforms integrate INO for semantic interaction classification, VO for vaccine hierarchy traversal, HDO for disease annotation, and DrugBank for drug association analysis, thereby enabling cross-entity biomedical exploration that surpasses conventional static interaction databases. Demonstrated use cases include ontology-guided vaccine network discovery, BioBERT-assisted interaction validation, AI-based literature summarization, and heterogeneous biomedical graph analyses. Furthermore, MCP-compatible interoperability enables external AI assistants to access biomedical interaction knowledge dynamically. Compared with PubTator, DisGeNET, KnetMiner, and related tools, Ignet 2.0 and Vignet uniquely combine daily PubMed literature updates, ontology-guided semantic analysis, BioBERT interaction scoring, and AI-assisted biomedical knowledge discovery within freely accessible web platforms.

## 6 Conclusion

Ignet 2.0 and Vignet function as ontology-guided platforms for biomedical literature mining and interaction analysis. They incorporate semantic interaction classification, heterogeneous biomedical network exploration, and artificial intelligence-assisted evidence interpretation. By integrating INO, VO, HDO, and DrugBank, these systems support standardized semantic annotation, vaccine-focused knowledge exploration, and cross-entity biomedical relationship analysis using continuously updated PubMed literature. Together, these capabilities enable scalable and transparent biomedical knowledge discovery within ontology-driven biomedical informatics and vaccine or drug ontology research.

## Acknowledgement

This platform is supported by the U.S. National Institute of Allergy and Infectious Disease (NIAID) [U24AI171008 to Y.H. and J.H.]

## Availability and requirements

Ignet 2.0 and Vignet are freely accessible web-based biomedical knowledge platforms available at https://ignet.org/ignet/ and https://ignet.org/vignet/, respectively. The systems are platform-independent and accessible via modern web browsers, with no local installation required. Programmatic access is supported via a public REST API (https://ignet.org/api/v1) and an MCP endpoint (https://ignet.org/api/v1/mcp) compatible with AI assistants. API endpoints are publicly accessible without registration. The database is updated daily from PubMed, and the source code is openly available at https://github.com/hurlab/Ignet.

